# CaMKK2 Identifies Biologically Aggressive Chronic Lymphocytic Leukemia and Regulates Leukemic Survival and Nurse-Like Cell Support

**DOI:** 10.64898/2026.02.23.707451

**Authors:** Shekeab Jauhari, Alicia D. Cooper-Volkheimer, Vini Verma, Dilber Gökçe Kaplan, Fahmin Basher, J. Brice Weinberg, Nelson Chao, Luigi Racioppi

## Abstract

**Background/Objectives:** Identification of prognostic biomarkers that capture biologically aggressive disease remains a major need in chronic lymphocytic leukemia (CLL). Aberrant calcium signaling contributes to leukemic survival; however, the clinical relevance of Ca²⁺/calmodulin-dependent protein kinase kinase 2 (CaMKK2), a calcium-responsive kinase, has not been defined. This study evaluated CaMKK2 as a candidate prognostic biomarker and functional regulator in CLL.

**Methods:** CaMKK2 expression was quantified in purified CD19⁺ CLL cells from a clinically annotated cohort balanced by immunoglobulin heavy chain variable region (IGHV) mutation status. Associations with time-to-treatment and overall survival were analyzed. Functional relevance was assessed by pharmacologic inhibition of CaMKK2 in primary CLL cells using metabolic (MTS) and apoptosis (Annexin V/PI) assays. Correlations between CaMKK2 expression and inhibitor sensitivity were determined. The impact of CaMKK2 inhibition on nurse-like cell (NLC) differentiation and macrophage-mediated leukemic support was evaluated in ex vivo culture systems.

**Results:** Elevated CaMKK2 expression was enriched in IGHV-unmutated CLL and associated with shorter time-to-treatment and inferior overall survival. CaMKK2 inhibition reduced primary CLL viability in a dose-dependent manner and induced apoptosis, with sensitivity correlating with CaMKK2 expression levels. Inhibition also attenuated CD163⁺ macrophage polarization and impaired NLC-mediated support of leukemic cells.

**Conclusions:** CaMKK2 expression identifies biologically aggressive CLL and functionally contributes to leukemic persistence. These findings position CaMKK2 as a prognostically relevant biomarker with therapeutic implications, supporting further evaluation of CaMKK2-targeted strategies in high-risk CLL.

**Sample Summary:** Chronic lymphocytic leukemia (CLL) shows marked variability in clinical outcome, highlighting the need for biomarkers that identify patients at higher risk of progression and guide therapeutic strategies. Calcium signaling supports leukemia cell survival, yet the clinical relevance of the calcium-responsive enzyme CaMKK2 has not been established. In this study, we demonstrate that elevated CaMKK2 expression in patient-derived leukemia cells is associated with more aggressive disease and earlier need for treatment. Laboratory experiments further show that inhibiting CaMKK2 reduces leukemia cell survival and disrupts supportive macrophage-like cells within the tumor microenvironment. These results position CaMKK2 as a candidate prognostic biomarker that reflects biologically high-risk disease and may inform therapeutic development. Future studies are warranted to determine whether CaMKK2-based risk stratification or targeted inhibition could improve management of patients with CLL.

## INTRODUCTION

Chronic lymphocytic leukemia (CLL) is a biologically heterogeneous B-cell malignancy characterized by progressive accumulation of CD5⁺CD19⁺ B cells in blood and lymphoid tissues. Clinical behavior ranges from indolent disease to rapidly progressive forms requiring early treatment[1,2]. Among established prognostic features, immunoglobulin heavy chain variable region (IGHV) mutation status remains a central determinant of outcome, with IGHV-unmutated CLL associated with enhanced B-cell receptor (BCR) signaling and inferior clinical prognosis[3,4].

CLL progression reflects both intrinsic leukemic cell programs and microenvironmental support. Within lymph node and marrow niches, stromal elements, T cells, and tumor-associated macrophages, including nurse-like cells (NLCs)[5,6], deliver cytokines, chemokines, and direct contact signals that reinforce BCR-associated pathways, sustain metabolic fitness, and limit apoptosis[7]. These niche-derived signals can attenuate responses to targeted therapies and contribute to minimal residual disease[8–10]. Thus, effective therapeutic strategies may require disruption of both leukemic survival pathways and macrophage-mediated niche protection.

Aberrant calcium signaling is a defining feature of CLL biology[11]. BCR activation promotes sustained intracellular Ca²⁺ flux through store-operated calcium entry (SOCE), mediated by ORAI1 and STIM channels[12]. Calcium-dependent signaling supports NFAT activation, metabolic adaptation, and apoptosis resistance[13]. While proximal BCR signaling has been successfully targeted with BTK inhibitors, emerging evidence suggests that downstream calcium-dependent programs may persist despite inhibition of upstream nodes[14]. The molecular effectors that couple SOCE to metabolic resilience in CLL remain incompletely defined.

Ca²⁺/calmodulin-dependent protein kinase kinase 2 (CaMKK2) is a serine/threonine kinase activated by Ca²⁺/calmodulin that regulates AMPK and related metabolic stress-response pathways[15]. CaMKK2 has been implicated in multiple malignancies, where it promotes tumor cell survival, macrophage polarization, and tumor progression[16–22]. In myeloid cells, CaMKK2 contributes to tumor-associated macrophage differentiation and immunosuppressive programming[23,24]. Recent work further links CaMKK2 to matrix-mediated mechanosensory signaling and AKT activation, positioning it at the convergence of calcium signaling and biomechanical adaptation[25]. However, the role of CaMKK2 in CLL, particularly in coordinating leukemic survival with macrophage-mediated niche support, has not been defined.

We hypothesized that CaMKK2 functions as a calcium-responsive signaling hub that integrates intrinsic leukemic survival programs with macrophage-mediated microenvironmental protection. To test this, we examined CaMKK2 expression in clinically annotated CLL cohorts, evaluated its association with disease progression, and assessed the impact of pharmacologic inhibition on primary CLL cells and NLC differentiation. Our findings identify CaMKK2 as a dual-compartment regulator of CLL persistence and a potential therapeutic target at the interface of leukemic fitness and niche support.

## MATERIALS AND METHODS

### CLL patient blood samples

Peripheral blood samples were obtained from CLL patients under Institutional Review Board-approved protocols Pro00011267at Duke University and the Durham VA Medical Center.

Patients were either treatment-naïve or had not received therapy within two years prior to peripheral blood collection. Available clinical and laboratory variables included age, sex, date of diagnosis, Rai stage, treatment history, cytogenetic abnormalities assessed by fluorescence in situ hybridization (FISH), and molecular prognostic markers, including IGHV mutation status, ZAP-70 expression, and CD38 expression. Clinical and laboratory characteristics of the patient cohorts are summarized in Supplemental Tables 1 and 3, and in the corresponding figure legends.

For analyses of CaMKK2 expression and clinical outcomes, a cohort of 40 patients was assembled and intentionally balanced by IGHV mutation status (20 IGHV-mutated and 20 IGHV-unmutated cases). IGHV status represents a major biological and prognostic determinant in CLL, reflecting differences in B-cell receptor signaling strength and disease aggressiveness[26]. This balanced design minimized subtype-driven bias, enabled direct comparison between risk groups, and permitted evaluation of CaMKK2 associations independent of cohort imbalance.

B lymphocytes were isolated from whole blood by negative selection using the RosetteSep™ Human B Cell Enrichment Cocktail (STEMCELL Technologies) and cryopreserved at -80 °C for RNA isolation and quantitative RT-PCR (qRT-PCR) analysis, as described in the corresponding section.

### Generation of nurse-like cells (NLCs)

NLCs were generated from CLL Peripheral blood mononuclear cells (PBMCs) as previously described[6]. Briefly, PBMC were isolated from DMSO-cryopreserved CLL PBMCs by density-gradient centrifugation using Ficoll-Paque (Sigma-Aldrich). PBMCs were resuspended in RPMI 1640 supplemented with 15% fetal calf serum and 1% penicillin-streptomycin and glutamine. PBMCs were then plated at high density (2 × 10^7^ cells/mL) in tissue culture-treated plates (Corning) and maintained at 37 °C in a humidified 5% CO_2_ atmosphere to allow differentiation of adherent NLCs. Where indicated, cultures were treated at the time of plating (day 1) with the CaMKK2 inhibitor STO-609 (5 µM; Tocris) or an equal volume of vehicle (Veh; DMSO). Supplemental doses of STO-609 (2.5 µM) or vehicle were added on days 5 and 10. To minimize drug accumulation while preserving early adherent precursors, partial medium changes were performed during treatment. After 14 days, non-adherent CLL-enriched cells were collected for analysis, and adherent cells were extensively washed with PBS to remove residual non-adherent cells. Adherent NLCs were then imaged directly by optical microscopy and subsequently detached using Macrophage Detachment Solution (PromoCell), according to the manufacturer’s instructions, and used for flow-cytometric phenotyping, or processed for RNA isolation and qRT-PCR, as indicated.

### Cell lines and preparation of Tumor-Conditioned Medium (TCM)

To model B-cell malignancy-derived factors within the tumor microenvironment, we used tumor-conditioned medium from human B cell tumor cell lines including OPM2, a multiple myeloma cell line derived from a patient with plasma cell leukemia and is commonly used as a model of aggressive plasma cell malignancy[27]; BJAB from Burkitt lymphoma[28], and SU-DHL-4, a diffuse large B-cell lymphoma originally derived from the peritoneal effusion of a 38-year-old male with non-Hodgkin lymphoma[29]. Briefly, tumor cells growing in logarithmic growth phase with viability >95% were harvested, washed twice with PBS, and resuspended in fresh complete medium at a density of 1-2 × 10^6^ cells/mL. Cultures were incubated at 37°C in a humidified 5% CO_2_ atmosphere for an additional 24-48h. Cell supernatants were then collected, centrifuged to remove cells, filtered through a 0.22µm membrane, aliquoted, and stored at -80 °C until use.

### Generation of Monocyte-Derived Macrophages (MDM) from healthy donor leukocytes

Leukocytes from healthy donors were obtained from the Gulf Coast Regional Blood Center as de-identified commercial human blood samples. Donor leukocytes were derived from single units of whole blood by density-gradient centrifugation. Donors were ≥16 years of age and met standard eligibility criteria for blood donation. All identifying information was removed prior to sample transfer, and samples were approved for in vitro research use; therefore, this protocol was determined to be IRB-exempt. Monocyte-derived macrophages were generated using a modified version of a previously described protocol[30]. Briefly, PBMC were resuspended in Monocyte Attachment Medium (PromoCell) and seeded into 12-well tissue culture plates at a density of 2-4 × 10^6^ cells per well. Cells were incubated at 37°C in a humidified 5% CO_2_ atmosphere for 1-2 h to allow monocyte adherence. Non-adherent cells were removed by extensive washing and adherent monocyte-enriched cells were cultured in macrophage differentiation medium (RPMI 1640 supplemented with recombinant human M-CSF 20 ng/mL; PeproTech), in the absence or presence of tumor-conditioned media (TCM; 50% v/v). STO-609 (5 µM) or an equal volume of vehicle (DMSO) was added at the time of plating (day 1). On day 3, culture media were partially replaced with fresh macrophage differentiation medium containing TCM or regular medium, with STO-609 or vehicle, corresponding to initial treatment conditions. After 6 days of differentiation, cultures were extensively washed with PBS, and adherent MDM were detached as described for NLC preparation. MDM immunophenotype was assessed by flow cytometry, and aliquots of cells were processed for quantitative RT-PCR analysis.

### MTS and Annexin V/PI assays

Primary CD19⁺ CLL cells (0.25 x 10^6^/well) were cultured in 96-well/plate in 0.1 ml of Hybridoma SFM (Gibco), as described[31–33]. Cells were treated with increasing concentrations of the CaMKK2 inhibitors STO-609[34] (Tocris), CC-8977[35] (Small Molecule Synthesis Facility Duke University), SGC-CaMKK2-1 (Sigma)[36], or an equal volume of vehicle (DMSO). Cells were then incubated at 37°C in a humidified 5% CO_2_ atmosphere, and CLL viability was assessed at 72 hours using two orthogonal approaches. Cytotoxicity assays were performed using the MTS [3-(4, 5-dimethylthiazol-2-yl)-5-(3-carboxymethoxyphenyl)-2-(4-sulfophenyl)-2H-tetrazolium] assay (CellTiter 96 Aqueous One Solution Cell Proliferation Assay; Promega), as described[37]. Briefly, MTS reagent was added to cell culture media and incubated for four hours before cell lysis by 1% SDS. Absorbance at 490 nm was measured using a microplate reader. The percentage of viable cells was determined by comparing the absorbance in drug-treated cells to the absorbance in vehicle-treated cells and calculate EC50 values. In parallel, apoptosis/cell death was quantified by flow cytometry using Annexin V-FITC (BD Biosciences) and propidium iodide (PI; Sigma) staining; total Annexin V⁺ events (PI⁻ and PI⁺ populations) were quantified across the same concentration range.

### Bright-field microscopy and ImageJ particle analysis

After 14 days of CLL-PBMC culture, non-adherent cells were removed, and wells were extensively washed with PBS to eliminate residual suspension cells. Adherent cells were imaged by bright-field optical microscopy using an Axiovert 200 microscope (Carl Zeiss Microscopy, Thornwood, NY) and identical acquisition settings across experimental conditions. For quantitative analysis, images were imported into ImageJ (NIH), and adherent structures were segmented using a consistent preprocessing and thresholding workflow applied uniformly across all images within an experiment. Particle abundance was quantified using the ImageJ Analyze Particles function and reported as particle number per unit area (particles/mm²).

### NLC-CLL co-culture assay

NLC from CLL PBMC were generated in the presence of vehicle or STO-609 (NLC-Veh and NLC-STO, respectively) as described above. After 14 days, CD19+ CLL were purified from the vehicle treated groups. purified. NLC-adherent cells from Vehicle and STO-609 groups were extensively washed, detached, and dispensed in 48-well plates at 1 × 10⁵ cells/well. CD19⁺ CLL cells (0.25 x 10^5^/well) were then cultured either alone (“None”) or co-cultured with autologous NLC-Veh or NLC-STO. At the indicated time points, non-adherent cells were collected, stained with CD19, and cell viability was quantified by flow cytometry as live CD19⁺ events

### Flow cytometry

Cells were harvested at the indicated time points, washed in PBS containing 2% FCS, and stained with fluorochrome-conjugated antibodies for surface markers, with inclusion of a fixable viability dye to exclude dead cells. All antibodies were used following manufacturer’s instructions. To assess CLL yield and viability, non-adherent cells were collected and stained with PE anti-human CD19 (clone HIB19, BioLegend) and live fixable dye (Zombie Near-IR; BioLegend). Adherent cells (NLC or MDM) were gently detached from wells using Monocyte Detachment Media (PromoCell), stained with Ig blocker, followed by saturating amounts of the following antibodies for 30 minutes at 4°C: anti-human AF488 anti-human HLA-DR (clone L234, BioLegend), PerCP/Cy5.5 anti-human/mouse CD11b (clone M1/70; BioLegend) or PE anti-human CD14 (clone HCD14, BioLegend) and live fixable dye (Zombie Near-IR; or Zombie yellow, BioLegend). Cells were fixed in 4% paraformaldehyde (PFA, Sigma), permeabilized with Tween-20, re-stained with PE/Cy7 anti-human CD68 (clone eBioY1/82A, eBioScience) and APC anti-human CD163 (clone GHI/61 eBioScience). All antibodies and staining procedures were performed according with manufacturer’s instructions. Unstained and single-stained controls were used for instrument setup and compensation, and fluorescence-minus-one (FMO) controls (with isotype controls as needed) were used to define gating thresholds. Cells were acquired with a BD FACSCanto cytometer and analyzed using FloJo 10.10.0 software.

### Gene expression analysis

Following PBMC culture, wells were washed and adherent cells were imaged as described above. Adherent cells were lysed directly in Buffer RLT (Qiagen) and collected by gentle scraping. Lysates were homogenized using QIAshredder columns (Qiagen) and total RNA was purified using the RNeasy Mini Kit (Qiagen) according to the manufacturer’s instructions. cDNA was synthesized from purified RNA using the SensiFAST cDNA Synthesis Kit (BIOLINE). Quantitative RT-PCR was performed using SYBR Green master mix, gene-specific forward and reverse primers (Integrated DNA Technologies), and cDNA template on an Applied Biosystems™ QuantStudio™ 6 Flex Real-Time PCR System. Primers sequences are listed in the Supplementary Table 4.

### Statistical analysis

Statistical analyses were performed using GraphPad Prism (version 10.6.1). Normality was assessed using Shapiro-Wilk testing and data distribution was not assumed to be normal. Overall survival and time-to-treatment were analyzed by Kaplan-Meier methods and compared using the log-rank (Mantel-Cox) test. For the independent validation cohort, survival analysis was performed using the SurvExpress platform[38]. For primary CLL drug-sensitivity assays, MTT dose-response curves were fit by nonlinear regression to derive EC50 values. Associations between CaMKK2 expression and STO-609 potency, and between metabolic inhibition and Annexin V-defined cell death, were assessed using Spearman rank correlation. For paired analyses of matched patient samples, statistical significance was assessed using two-tailed paired tests. For co-culture experiments with multiple conditions and time points, statistical significance was determined by two-way ANOVA with appropriate multiple-comparisons correction. For tumor-conditioned media experiments, differences across conditions were evaluated by ANOVA with multiple-comparisons testing, including planned comparisons of vehicle versus STO-609 within each conditioned-media condition. Unless otherwise stated, all tests were two-sided.

## RESULTS

### Elevated *CAMKK2* expression associates with high-risk CLL biology and clinical course

To evaluate CaMKK2 as a candidate biomarker in CLL in the context of established prognostic factors, we analyzed clinically annotated CLL biorepository specimens available at the Division of Hematological Malignancies and Cellular Therapy (Duke University). We selected 40 cryopreserved peripheral blood CLL samples, balanced a priori by IGHV mutation status (20 IGHV-mutated; 20 IGHV-unmutated), and quantified *CaMKK2* mRNA in purified CD19⁺ CLL cells by qRT-PCR. Cohort characteristics are summarized in **Supplementary Table 1**. Because CaMKK2 quantification required adequate RNA and complete clinical linkage, *CaMKK2* expression was evaluable in 33 samples. Patients were stratified using the cohort median of *CaMKK2* expression (high versus low; group sizes shown in the plots), a pragmatic and unbiased threshold that enables balanced comparison without data-driven optimization. Using this cutoff, CaMKK2-high status was associated with significantly shorter time-to-treatment (log-rank p = 0.0001; **Figure 1A**) and inferior overall survival (log-rank p = 0.0385; **Figure 1B**), supporting its potential clinical relevance as a biomarker of adverse outcome. *CaMKK2* expression, normalized by *ACTB*, was significantly higher in IGHV-unmutated compared with IGHV-mutated CLL (**Figure 1C**). Further, the Duke cohort recapitulated canonical IGHV-linked biology, with higher CD38 and ZAP-70 in IGHV-unmutated cases (**Supplementary Figure S1**). In an independent dataset (GSE22762; Herold et al.[39]), *CaMKK2*-high expression showed a directionally consistent adverse association with survival (SurvExpress[38]; **Figure 1D**). Finally, in a multivariable model including IGHV status and Rai stage, IGHV-unmutated status remained the dominant correlate of higher *CaMKK2* expression (β = 0.496, 95% CI 0.310-0.682, p = 1.68 × 10⁻^7^), whereas Rai stage was not independently associated (p = 0.307; **Supplementary Table S2**). Collectively, these data show that *CaMKK2* expression in blood CLL cells is enriched in the IGHV-unmutated subtype and associates with a more aggressive clinical course.

**Figure 1.**
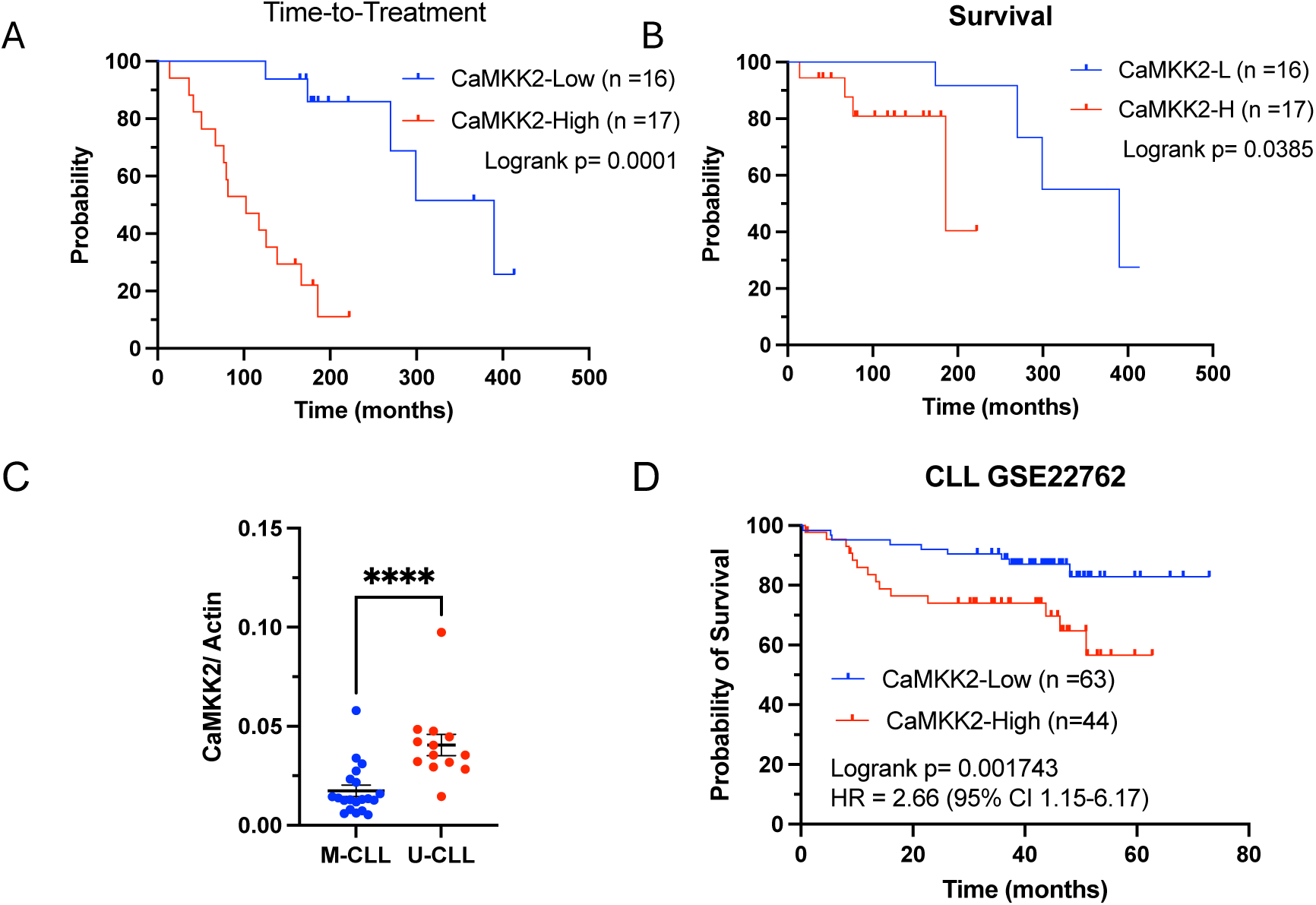
CaMKK2 expression in CLL associates with adverse outcome and aggressive disease biology. (A,B) Kaplan-Meier analyses of time-to-treatment (A) and overall survival (B) for CLL patients stratified by *CaMKK2* expression using a median split (CaMKK2-Low, n = 16; CaMKK2-High, n = 17). Tick marks indicate censored observations. P values were calculated using the log-rank (Mantel-Cox) test. (C) CaMKK2 mRNA expression in CD19⁺ CLL cells obtained from the Duke CLL biorepository, quantified by qRT-PCR and normalized to ACTB. Samples are grouped by IGHV mutation status (IGHV-mutated [M-CLL] vs IGHV-unmutated [U-CLL]). Each symbol represents one patient; horizontal bars indicate mean ± SEM. Statistical significance was assessed by two-tailed Mann–Whitney test (**** p < 0.0001). (D) Independent cohort validation using the CLL dataset reported by Herold et al. (GSE22762). Overall survival was analyzed using the SurvExpress platform, stratifying cases into CaMKK2-high versus CaMKK2-low groups. Log-rank p value and hazard ratio (HR) with 95% confidence interval (CI) are indicated.

### Pharmacologic CaMKK2 inhibition with STO-609 reduced primary CLL viability and induced Annexin V-defined cell death in a dose-dependent manner

To test whether CaMKK2 represents a therapeutic vulnerability in CLL, purified primary CD19⁺ CLL cells from nine independent patient samples (clinically annotated in **Supplementary Table S3**) were treated with increasing concentrations of the CaMKK2 inhibitor STO-609 and evaluated using orthogonal assays of metabolic activity (MTS) and cell death (Annexin V/PI) (**Figure 2**). Representative dose-response curves from three samples (IDs 824-003, 400-013, and 558-013) demonstrated a concentration-dependent reduction in MTS signal accompanied by an increase in total Annexin V⁺ events (including PI⁻ and PI⁺ populations) (**Figure 2A,B**). Across the nine samples, STO-609 sensitivity varied (MTS-derived EC50 range ∼0.97-7.67 µM, providing clinically annotated context for inter-sample heterogeneity in response. Consistent with on-target biology, STO-609 potency correlated with CaMKK2 expression measured by qRT-PCR (*CaMKK2*/*ACTB*), with higher *CaMKK2* associated with greater sensitivity (Spearman r = -0.85, p = 0.0061; **Figure 2C**). To distinguish STO-609-induced cytotoxicity from isolated metabolic suppression, we compared MTS response and Annexin V response at a fixed dose. At 5 µM STO-609, MTS viability (% of vehicle control) inversely correlated with the increase in total Annexin V positivity (treated - vehicle; normalized to each sample’s maximal response at 20 µM) (Spearman r = -0.9333, p = 0.0007; **Figure 2D**). To further assess whether the observed effects were specific to STO-609, two additional structurally distinct CaMKK2 inhibitors, SGC- CaMKK2-1[36] and CC-8977[35], were evaluated in CLL primary cells from three independent patients, including one with documented multi-drug-resistant disease (**Supplementary Table S4,** and **annotated in Supplementary Table S3**). Both compounds reduced CLL survival in MTS assay, demonstrating activity broadly comparable to STO-609, although potency varied across samples. Notably, CC-8977 exhibited higher EC50 in the multi-drug-resistant sample. These findings provide supportive evidence functional evidence that inhibition of CaMKK2 activity compromises the survival of primary CLL cells beyond a single inhibitor scaffold. The integration of clinical associations and pharmacologic data supports CaMKK2 as both a biomarker of adverse disease biology and a tractable therapeutic target in high-risk CLL.

**Figure 2.**
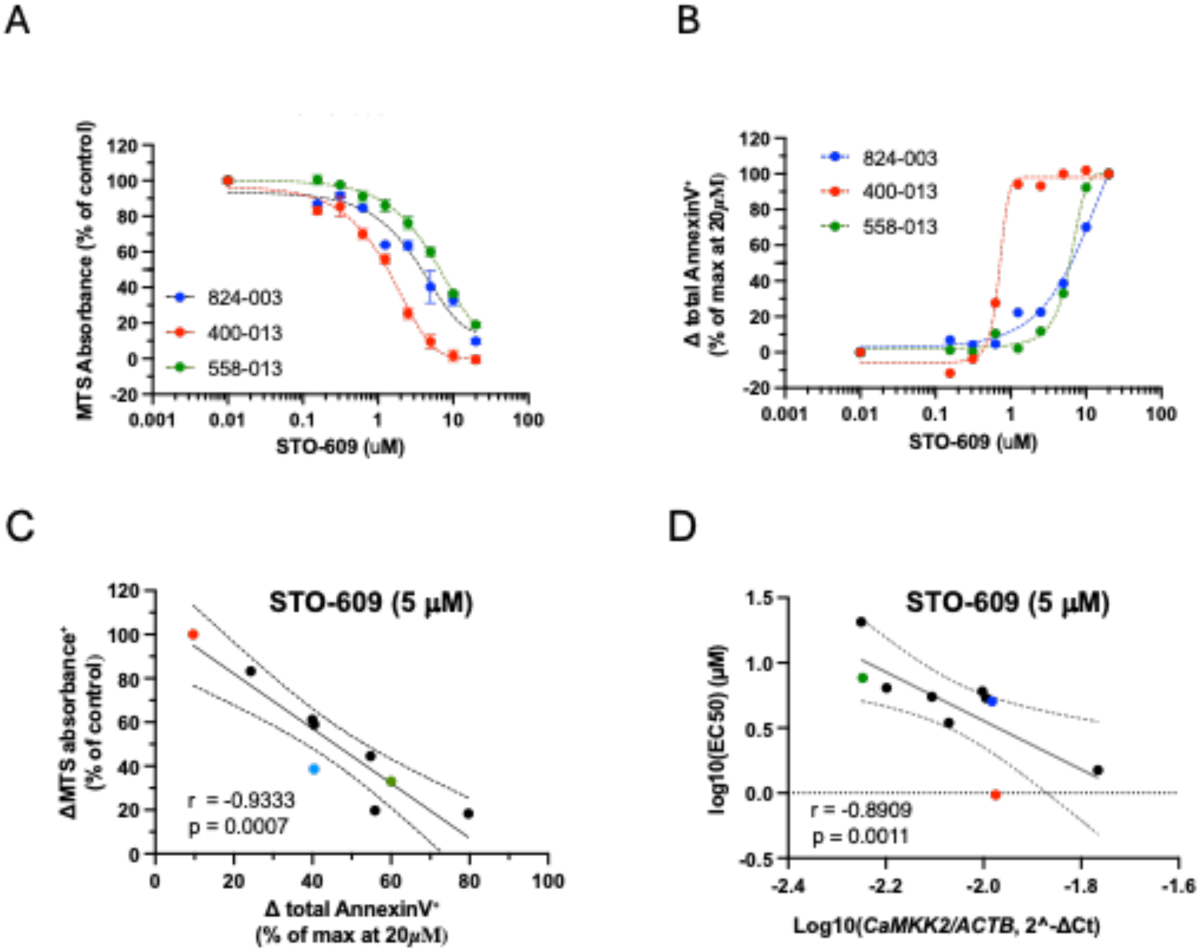
CaMKK2 inhibition reduces viability and induces apoptosis in primary CLL cells. Primary CD19⁺ purified CLL cells (n = 9 independent patient samples) were treated with increasing concentrations of the CaMKK2 inhibitor STO-609 and assessed using complementary assays of metabolic activity and cell death. (A) Representative MTS dose-response curves from three CLL samples (IDs 824-003, 400-013, and 558-013); absorbance is expressed as percentage of vehicle control. (B) Representative flow-cytometry dose-response curves from the same samples, reporting total Annexin V⁺ cells (PI⁻ and PI⁺ populations). (C) Relationship between metabolic inhibition and cell death at 5 µM STO-609, plotting MTS (% of control) versus Δ total Annexin V⁺ (treated - vehicle), normalized to the maximal response observed at 20 µM for each sample. (D) Association between STO-609 potency and *CaMKK2* expression across primary CLL samples (n = 9), plotting MTS-derived EC50 values versus *CaMKK2* expression measured by qRT-PCR and normalized to *ACTB*. Associations were assessed using Spearman rank correlation; regression lines are shown for visualization.

### CaMKK2 inhibition during PBMC culture reduces CLL cell viability and alters NLC yield and phenotype

CLL persistence is supported by both cell-autonomous survival programs within the leukemia cell and cell-extrinsic cues provided by the myeloid microenvironment. NLCs provide an established *ex vivo* model of this protective niche[6], while *CaMKK2* is highly expressed in pro-tumoral macrophage states[23,25,40], raising the possibility that CaMKK2 blockade could disrupt the CLL-NLC cross-functional axis that sustains leukemic survival. To test this hypothesis, PBMCs from CLL patients were cultured at high density for 14 days in the presence of vehicle or STO-609 (5 µM) to generate adherent NLC (**Figure 3**; and annotated in **Supplementary Table 3**). Non-adherent and adherent cells were recovered as described in Material and Methods section and analyzed for yield, phenotype, and gene expression. Compared with vehicle, STO-609 treatment significantly reduced the percentage of viable CD19⁺ CLL cells recovered in the non-adherent fraction across paired patient samples (**Figure 3A-B**). Bright-field imaging after extensive washing of well plates showed reduced adherent cell yield with STO-609, which was confirmed by ImageJ-based particle quantification (**Figure 3B**). Phenotypic analysis of detached adherent cells demonstrated a decrease in the proportion of CD163⁺ cells within the CD68⁺ macrophage compartment (Figure 3C), consistent with impaired acquisition of an NLC-like phenotype (**Figure 3C**). At the transcript level, *CaMKK2* was detectable in CLL and, in paired samples, was expressed at higher levels in NLC than in autologous CLL cells (**Supplementary Figure S2**). STO-609 treatment altered expression of macrophage-associated inflammatory mediators (*IL6* and *CXCL10*)[41–44], which are positively associated with leukemia cell survival and disease progression, whereas *CaMKK2*, *BAFF*, and *APRIL* mRNA levels in adherent NLC were not significantly changed (**Figure 3D**). Collectively, these data show that CaMKK2 blockade during PBMC culture reduces CLL cell persistence while limiting the formation of an adherent, CD163⁺ NLC compartment, supporting CaMKK2 as a modifiable cross-cell pathway required for establishment of a pro-tumoral, protective niche in CLL.

**Figure 3.**
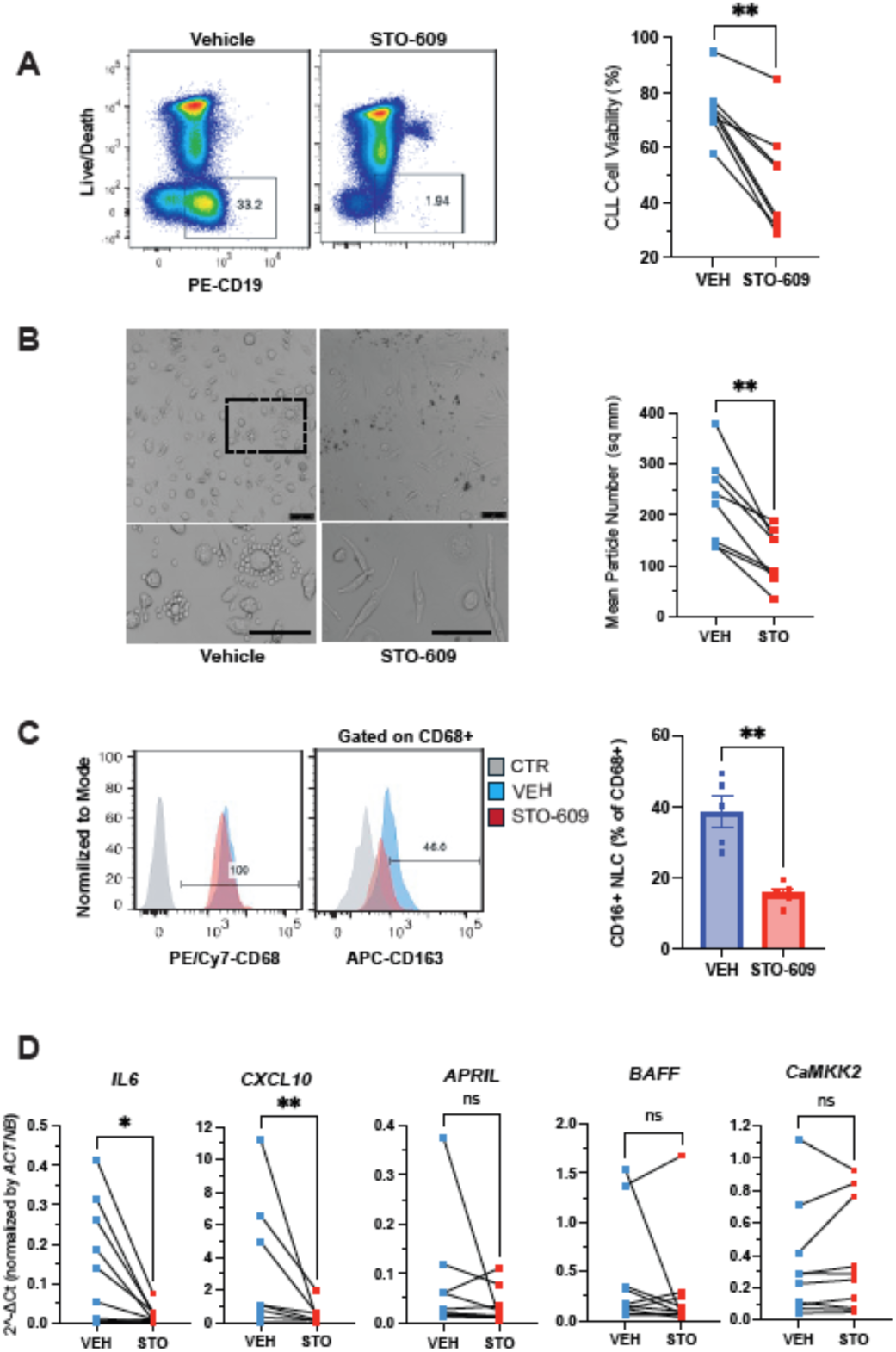
CaMKK2 inhibition during PBMC culture reduces leukemia cells viability and nurse-like cells (NLC) generation. Peripheral blood mononuclear cells (PBMCs) from CLL patients were cultured at high density for 14 days in the presence of vehicle (Veh; DMSO) or the CaMKK2 inhibitor STO-609 (5 µM). Non-adherent and adherent fractions were analyzed separately. (A) Non-adherent cells were stained for CD19 and viability markers, and CD19⁺ CLL cell viability was quantified by flow cytometry. Representative plots are displayed on the left. Paired data from individual patients are shown on the right. (B) Adherent cells were visualized by bright-field microscopy after extensive washing. (Left) Representative images are shown; scale bar = 75 µm. Insets indicate regions shown at higher magnification. Images were analyzed using ImageJ, and the mean number of adherent particles per mm² is quantified (Right). (C) Adherent cells were detached and analyzed by flow cytometry for macrophage markers. Representative histograms of CD68 and CD163 expression are shown. The right panel quantifies the percentage of CD163⁺ cells within the CD68⁺ population. (D) RNA isolated from adherent cells was analyzed by qRT-PCR to quantify expression of genes of interest normalized to *ACTNB* and expressed as 2⁻ΔCt. Each line represents paired measurements from an individual patient. Statistical significance was assessed using paired two-tailed tests; *, **, and *** indicate p < 0.05, p < 0.01, and p < 0.001, respectively.

### STO-609 Impairs NLC-Mediated Support of CLL Cell Survival

Building on our findings that CaMKK2 expression stratifies CLL clinical behavior (**Figure 1**; **Supplemental Table S1**) and that pharmacologic CaMKK2 inhibition directly compromises primary CLL viability (**Figure 2**), we next evaluated whether CaMKK2 blockade also weakens CLL survival under conditions of myeloid microenvironmental support. PBMCs from CLL patients were cultured at high density for 14 days with vehicle or STO-609 (5 µM) to generate adherent NLC, after which NLC from vehicle- or STO-609-treated cultures were detached, replated, and used as feeder cells for autologous CD19⁺ CLL cells purified from the vehicle condition (**Figure 4**, and annotated **in Supplementary Table 3**). As expected, CLL cells cultured alone (“None”) showed progressive loss of viability over time, whereas co-culture with vehicle-derived NLC (NLC-Veh) increased recovery of live non-adherent CD19⁺ cells at days 3 and 7, confirming the protective function of NLC in this system (**Figure 4A,B**). In contrast, NLC generated in the presence of STO-609 (NLC-STO) exhibited a markedly reduced capacity to sustain CLL viability, resulting in significantly fewer live CD19⁺ cells compared with NLC-Veh at both time points (**Figure 4A,B**). These effects were reproducible across three independent CLL patient samples, and two-way ANOVA (time × feeder condition) with multiple-comparison correction, supporting a significant impact of NLC conditioning on CLL survival (**Figure 4C**). Together, these data demonstrate that CaMKK2 blockade impairs the pro-survival function of NLC, disrupting CLL-NLC crosstalk and weakening formation of a macrophage-dependent protective niche that sustains leukemic cell persistence.

**Figure 4.**
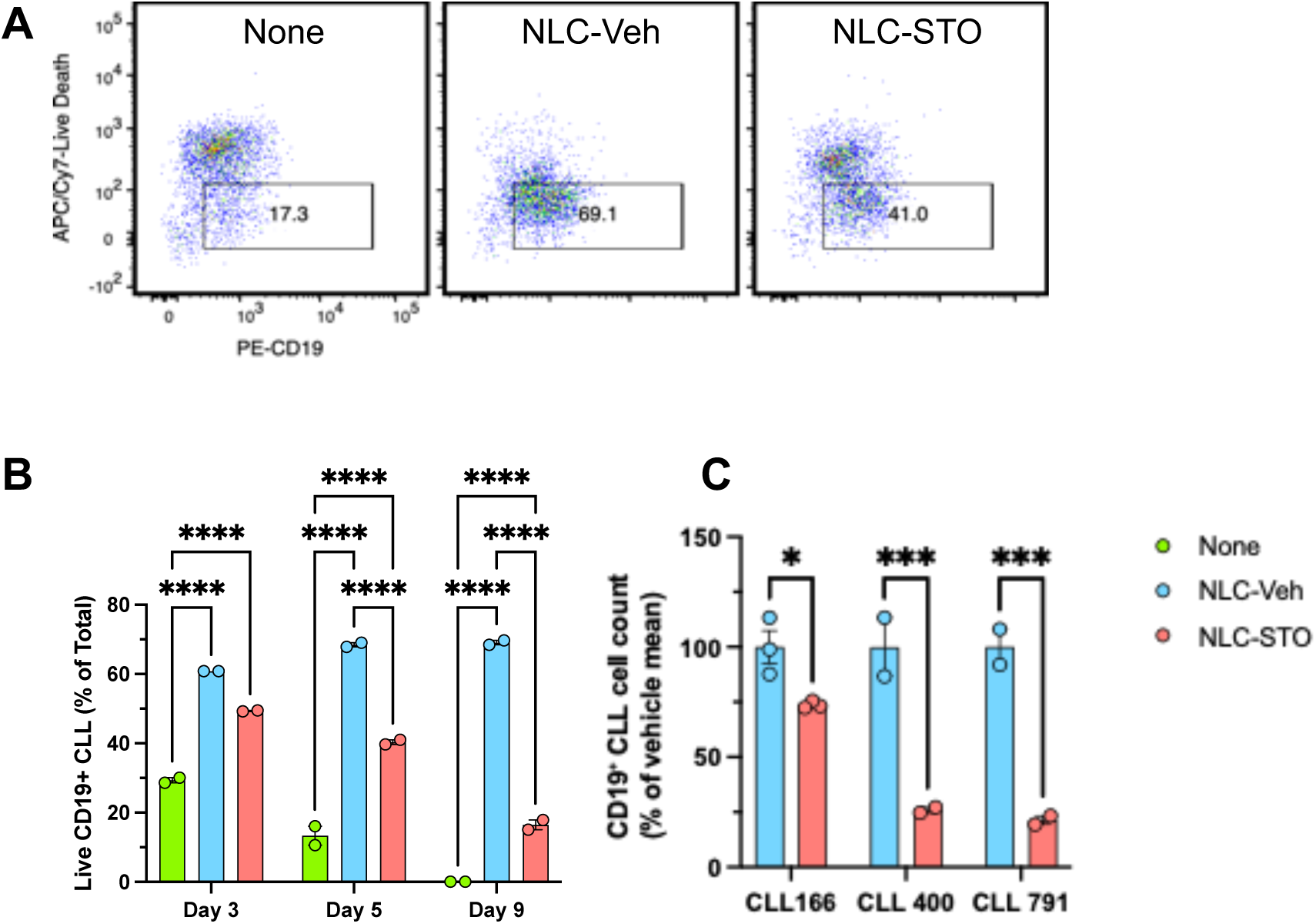
Blocking CaMKK2 during PBMC culture impairs NLC-mediated support of CLL cell survival. PBMCs from CLL patients were cultured at high density for 14 days with vehicle or STO-609 (5 µM) to generate adherent NLC. CD19⁺ CLL cells were purified from vehicle cultures, while NLC from vehicle- or STO-609-treated cultures were detached and replated. Purified CLL cells were cultured alone (“None”) or co-cultured with NLC generated under vehicle or STO-609 conditions. (A) Representative flow-cytometry plots of viable CD19⁺ CLL cells at days 3 and 7 from a representative patient (CLL 791). (B) Quantification of live CD19⁺ CLL cells for CLL 791 at the indicated time points. (C) Summary of live CD19⁺ CLL cell percentages from three independent CLL patients. Data are shown as mean ± SEM. Statistical significance was determined by two-way ANOVA with multiple-comparison correction; **, ***, and **** indicate p < 0.01, p < 0.005, and p < 0.001, respectively.

### Tumor-conditioned media drives an immunosuppressive macrophage phenotype that is attenuated by CaMKK2 inhibition

To test whether CaMKK2 contributes to tumor-driven programming of macrophage phenotypes relevant to the CLL microenvironment, we generated monocyte-derived macrophages (MDM) from 3 independent healthy donor leukocytes and cultured them either in control conditions (“None”) or in the presence of tumor-conditioned media (TCM) from the B-cell malignancy cell lines OPM2, SH-DHL-4, or BJAB, with vehicle or STO-609 (5 µM) (**Figure 5**). Flow-cytometric analysis of CD68⁺ macrophages showed that exposure to TCM increased the emergence of a CD163⁺ subset (**Figure 5A-C**), consistent with induction of a pro-tumoral, NLC-like phenotype by tumor-derived signals. Across conditioned media sources, CaMKK2 blockade with STO-609 attenuated this response, reducing CD163 expression and/or the frequency of CD163⁺ macrophages relative to matched vehicle controls (Figure 5B,C), while preserving the overall CD68⁺ macrophage compartment (**Figure 5A,B**). Together with the patient-derived NLC data (**Figures 3-4**), these findings support a model in which CaMKK2 functions as a tractable signaling node linking tumor-derived cues to macrophage polarization and microenvironmental support, thereby providing a mechanistic basis for how CaMKK2 inhibition can disrupt pro-tumoral myeloid programming and weaken the protective niche that sustains CLL.

**Figure 5.**
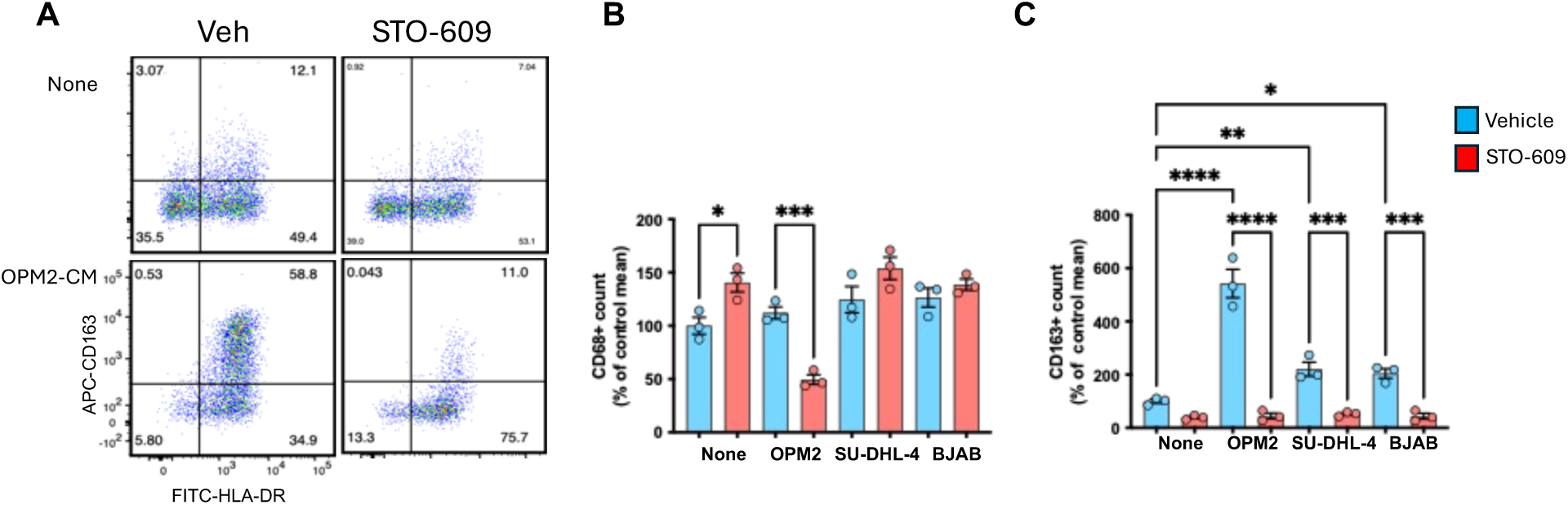
Tumor-derived signals induce an immunosuppressive NLC phenotype that is attenuated by CaMKK2 inhibition. Monocyte-derived macrophages from healthy donors were cultured in the absence (“None”) or presence of tumor-conditioned media, with vehicle or STO-609 (5 µM), and analyzed by flow cytometry. (A) Representative flow-cytometry plots showing HLA-DR and CD163 expression on CD14⁺CD68⁺ macrophages cultured with or without OPM2-conditioned medium (OPM2-CM). (B,C) Quantification of total CD68⁺ macrophages and the CD163⁺ subset following exposure to conditioned media from OPM2, SU-DHL-4, or BJAB cells. CaMKK2 inhibition limits acquisition of a CD163⁺ immunosuppressive phenotype induced by tumor-derived factors. Data represent paired analyses from independent experiments. Statistical significance was determined using paired tests; significance levels are indicated in the figure.

## DISCUSSION

In this study, we identify CaMKK2 as a calcium-responsive signaling node that operates across both leukemic and microenvironmental compartments in CLL. Our data support a model in which CaMKK2 integrates intrinsic leukemic survival pathways with macrophage-mediated niche support, thereby contributing to disease persistence and therapeutic resistance.

At the tumor cell level, elevated CaMKK2 expression in purified CD19⁺ CLL cells was associated with shorter time-to-treatment and inferior overall survival and was enriched in IGHV-unmutated disease. Given that IGHV-unmutated CLL exhibits heightened and sustained BCR signaling and enhanced calcium flux[3,4], these findings suggest that CaMKK2 marks a calcium-dependent, metabolically resilient leukemic state.

Mechanistically, store-operated calcium entry (SOCE) via ORAI1/STIM channels sustains intracellular Ca²⁺ flux that promotes NFAT activation and apoptosis resistance[12,45]. CaMKK2 functions downstream of Ca²⁺/calmodulin to activate AMPK metabolic stress-adaptation programs[15], providing a mechanistic link between calcium entry and mitochondrial fitness[24,46]. The positive correlation between CaMKK2 expression and STO-609 sensitivity reinforce the concept that CaMKK2 functions as a downstream amplifier of SOCE-mediated survival signaling. Notably, recent structural analyses of aggressive stereotyped CLL subset 1 indicated that autonomous BCR-BCR homotypic signaling is not universally conserved, suggesting that alternative mechanisms must sustain intracellular signaling and leukemic fitness. In this context, CaMKK2-dependent signaling may represent a parallel or compensatory pathway reinforcing disease progression. Unlike BTK inhibitors, which target proximal BCR signaling nodes, CaMKK2 inhibition disrupts calcium-driven metabolic resilience, identifying a mechanistically distinct vulnerability.

CLL progression depends on intrinsic biological features of leukemia cells as well as niche-mediated support[47,48]. Within lymph node and marrow niches, NLCs and other myeloid populations provide cytokines and direct contact signals that reinforce BCR-associated pathways, sustain metabolic adaptation, and limit apoptosis. These inputs can attenuate responses to targeted agents and contribute to minimal residual disease. Our data demonstrate that CaMKK2 inhibition disrupts this protective axis. STO-609 reduced the yield and of CD163⁺ macrophage, altered inflammatory mediator expression, and impaired macrophage-mediated survival of autologous CLL cells. These findings align with prior work showing that myeloid CaMKK2 regulates tumor-associated macrophage functions and promotes tumor progression in breast cancer and lymphoma models[23,24]. Importantly, recent evidence linking CaMKK2 to matrix-mediated mechanosensory signaling and AKT activation further expands this framework[25]. Lymph node remodeling and altered extracellular matrix composition are recognized features of aggressive CLL, and CaMKK2 may function at the convergence of calcium signaling, metabolic adaptation, and biomechanical cues within these niches[5,7–10]. Thus, CaMKK2 appears to coordinate biochemical and structural survival inputs across both leukemic and myeloid compartments.

The dual intrinsic-extrinsic activity of CaMKK2 carries important therapeutic implications. While BTK and BCL2 inhibitors disrupt key survival pathways, microenvironmental protection remains a barrier to durable disease control. Increasingly evidence indicates that macrophage-rich, metabolically competitive, and structural remodeled niches contribute to the comparatively limited durability of CAR-T cell therapy in CLL relative to other B-cell malignancies[49–51]. Immunosuppressive macrophages, cytokine gradients, and matrix-associated constraints can impair T-cell fitness and persistence. By simultaneously reducing leukemic metabolic resilience and attenuating macrophage-mediated support, CaMKK2 inhibition may recalibrate this hostile microenvironment. Although *in vivo* validation is required, our findings suggest that targeting CaMKK2 could enhance the efficacy of both targeted agents and immune-based therapies by disrupting the bidirectional crosstalk that sustains leukemic persistence.

Several limitations warrant consideration. STO-609 has been widely used as selective CaMKK2 inhibitor in pre-clinical models[23,24,52–56], and additional compounds such as SGC-CaMKK2-1 and CC-8977 have demonstrated activity and selectivity across various systems[35,36,57].

However, pharmacologic approaches, while translationally relevant, may have off-target effects. Future studies employing genetic perturbation strategies or next-generation clinical-grade inhibitors will be important to define the specific contribution of CaMKK2 to CLL biology. In addition, our microenvironmental experiments rely on *ex vivo* PBMC-derived models which capture key aspects of NLC biology but do not fully reproduce the spatial and cellular complexity of lymph node niches. Finally, while the cohort was intentionally balanced by IGHV status, validation in larger, independently annotated datasets with multivariable modeling will be necessary to confirm the prognostic utility of CaMKK2 in CLL.

## Conclusions

CaMKK2 emerges from this study as a clinically relevant biomarker and functional regulator of chronic lymphocytic leukemia. Elevated expression identifies biologically aggressive disease and associates with adverse clinical outcomes, while functional inhibition reduces leukemic cell viability and weakens macrophage-mediated microenvironmental support. These findings position CaMKK2 at a critical junction between tumor-intrinsic survival and niche protection. By linking prognostic relevance with actionable biology, CaMKK2 represents a promising candidate for biomarker-guided risk stratification and therapeutic development in high-risk CLL.

## Institutional Review Board Statement

The study was conducted in accordance with the Declaration of Helsinki and approved by the Institutional Review Board of Duke University and the Durham VA Medical Center (IRB protocol number Pro00011267).

## Informed Consent Statement

Informed consent was obtained from all subjects involved in the study.

## Data Availability Statement

The datasets generated and/or analyzed during the current study are available from the corresponding author upon reasonable request. Publicly available transcriptomic data used for validation were obtained from the Gene Expression Omnibus (GEO) under accession number GSE22762.

## Author Contributions

Conceptualization (SJ, LR, JBW, NC); methodology (SJ, LR, ADCV, FB); formal analysis (SJ, LR); writing original draft preparation (SJ, LR, JBW, VV, DJK, NC); All authors have read and agreed to the published version of the manuscript.

