## Supplementary files for "CaMKK2 Identifies Biologically Aggressive Chronic Lymphocytic Leukemia and Regulates Leukemic Survival and Nurse-Like Cell Support"

**By**

Shekeab Jauhari, Alicia D. Cooper-Volkheimer, Vini Verma, Dilber Gökçe Kaplan, Fahmin Basher, J. Brice Weinberg, Nelson Chao, and Luigi Racioppi.

**Supplementary Table S1. Demographic and clinical summary of CLL patient samples used for association studies (Figure 1).**

| | Overall | IGHV mutated | IGHV unmutated | CaMKK2 cohort |
|---|---|---|---|---|
| **N** | 40 | 20 | 20 | 33* |
| **Age at Dx, median [IQR]** | 60.0 [55.0-68.0] | 60.0 [56.8-68.0] | 59.0 [53.0-66.0] | 61.0 [57.0-68.0] |
| **Female** | 10 (25.0%) | 7 (35.0%) | 3 (15.0%) | 9 (27.3%) |
| **Male** | 30 (75.0%) | 13 (65.0%) | 17 (85.0%) | 24 (72.7%) |
| **Rai 0** | 26 (65.0%) | 19 (95.0%) | 7 (35.0%) | 24 (72.7%) |
| **Rai 1–2** | 11 (27.5%) | 1 (5.0%) | 10 (50.0%) | 6 (18.2%) |
| **Rai 3–4** | 3 (7.5%) | 0 (0.0%) | 3 (15.0%) | 3 (9.1%) |
| **IGHV unmutated** | 20 (50.0%) | 0 (0.0%) | 20 (100.0%) | 14 (42.4%) |
| **CD38 ≥30%** | 10 (25.0%) | 1 (5.0%) | 9 (45.0%) | 8 (24.2%) |
| **ZAP70 ≥20%** | 24 (60.0%) | 10 (50.0%) | 14 (70.0%) | 18 (54.5%) |
| **13q del (any)** | 20 (50.0%) | 12 (60.0%) | 8 (40.0%) | 16 (48.5%) |
| **11q del (any)** | 5 (12.5%) | 1 (5.0%) | 4 (20.0%) | 5 (15.2%) |
| **17p del (any)** | 2 (5.0%) | 1 (5.0%) | 1 (5.0%) | 2 (6.1%) |
| **trisomy 12 (any)** | 5 (12.5%) | 1 (5.0%) | 4 (20.0%) | 4 (12.1%) |
| **normal FISH** | 12 (30.0%) | 6 (30.0%) | 6 (30.0%) | 10 (30.3%) |
| **OS, months median [IQR]** | 159.8 [79.8-184.7] | 186.3 [174.3–230.7] | 79.3 [47.8-117.6] | 164.1 [100.8-195.2] |
| **TTT, months median [IQR]** | 87.4 [10.9-178.7] | 178.8 [168.5–211.3] | 10.7 [3.0-13.5] | 159.6 [12.8-179.1] |
| **Deaths** | 11 (27.5%) | 5 (25.0%) | 6 (30.0%) | 8 (24.2%) |

*Samples assessed for *CaMKK2* gene expression.

**Supplementary Table S2. Multivariable linear regression of CaMKK2 expression with IGHV mutation status and Rai stage (Duke CLL cohort).**

| Predictor | Beta_log10 | CI95-low | CI95-high | Fold change-10^beta | p |
|---|---|---|---|---|---|
| IGHV unmutated (vs mutated) | 0.4960 | 0.3102 | 0.6818 | 3.1330 | 1.6766e-07 |
| Rai stage, per 1 unit | -0.0565 | -0.1647 | 0.0518 | 0.8781 | 0.3068 |

**Supplementary Table S3. Demographic and clinical summary of CLL patient samples used for *in vitro* studies.**

| CLL ID# | TTT (Year) | Rai | IGVH Mut | CD38 | ZAP70 | FISH | FISH | Age at Dx | Sex | Fig 3A | Fig 2/TabS4 | Fig 3/Fig S2 | Fig 4 |
|---|---|---|---|---|---|---|---|---|---|---|---|---|---|
| 46 | 2.19 | 0 | - | - | + | 13q | good | 58 | female | V | | | |
| 69 | 15.6 | 0 | - | + | + | 13q | good | 62 | male | | V | | |
| 166 | N/A | 0 | + | - | + | normal | good | 44 | female | | | | V |
| 183 | 13.66 | 0 | + | - | - | 13q | good | 59 | female | V | | | |
| 276 | N/A | 0 | + | - | - | 13q | good | 61 | female | | | V | |
| 338 | N/A | 0 | - | - | + | 13q | good | 64 | male | | | V | |
| 400 | 2.53 | 0 | + | - | - | 13q | good | 48 | male | | V | | V |
| 467 | N/A | 0 | + | - | - | 13q | good | 71 | female | | | V | |
| 490 | 8.63 | 0 | + | - | - | normal | good | 65 | female | V | | | |
| 499 | 15.54 | 0 | + | - | + | N/A | N/A | 57 | male | | V | | |
| 558 | 5.25 | N/A | + | - | + | 13q | good | 47 | male | V | V | | |
| 569 | N/A | N/A | + | - | - | 13q | good | 59 | male | | | V | |
| 608 | 3.23 | N/A | - | - | - | normal | good | 63 | male | V | | | |
| 621 | 5.39 | 0 | N/A | - | - | 13q | good | 59 | male | | V | | |
| 626 | N/A | 0 | + | - | - | normal | good | 67 | female | V | | | |
| 631 | 5.3 | N/A | + | + | + | N/A | N/A | 63 | male | V | | | |
| 642 | 4.76 | 0 | - | - | + | 11q13q | bad | 68 | male | V | | | |
| 643 | 4.92 | 0 | - | N/A | N/A | 13q | good | 54 | male | | | V | |
| 647 | N/A | 0 | + | - | - | 13q | good | 48 | female | | V | | |
| 649 | 6.79 | | + | - | + | 13q | good | 53 | male | V | | V | |
| 673 | 3.66 | N/A | - | N/A | + | tri12 | ntermediate | 53 | male | | | V | |
| 686 | 3.97 | 0 | - | + | + | normal | good | 61 | female | | | V | |
| 700 | 2.45 | 0 | + | + | - | 13q | good | 69 | male | | V | | |
| 714 | 11.18 | 0 | + | N/A | N/A | 13q | good | 52 | male | | | V | |
| 716 | 4.62 | 0 | - | - | + | 13q | good | 59 | female | V | | | |
| 723 | N/A | 0 | + | - | - | 13q | good | 73 | male | | | V | |
| 729 | 1.8 | 0 | + | - | - | normal | good | 66 | male | | V | | |
| 733 | 0.08 | N/A | N/A | - | N/A | 17p13q | bad | 68 | female | | V | | |
| **763** | 4.48 | 0 | - | - | + | 13q | good | 64 | male | V | V | | |
| 766 | 4 | N/A | + | N/A | N/A | 11q13q | bad | 35 | male | | | V | |
| 768 | N/A | 0 | N/A | - | - | 13q | good | 41 | male | | | V | |
| 781 | N/A | N/A | + | N/A | N/A | 13q | good | 71 | male | | | V | |
| 783 | N/A | 0 | N/A | N/A | N/A | 13q | good | 56 | male | | | V | |
| 791 | N/A | 0 | N/A | N/A | N/A | tri12 | ntermediate | 56 | male | | | V | V |
| 824 | 0.16 | 0 | N/A | - | + | 11q13q | Bad | 54 | male | | V | | |
| 842 | N/A | 0 | + | - | - | 13q | good | 67 | male | V | | V | |
| 846 | 11.94 | 0 | + | N/A | N/A | tri12 | ntermediate | 62 | female | | | V | |
| 862 | N/A | 1 | - | + | N/A | normal | good | 73 | male | | | V | |
| 866 | 0.5 | 1 | N/A | - | + | normal | good | 66 | male | | V | | |

**Supplementary Table S4**

| | Patient | Patient | Patient |
|---|---|---|---|
| **Drug ED50 (uM)** | **621-007** | **647-011** | **733-012*** |
| SGC-CaMKK2-1 | 5.29 | 5.49 | 27.71 |
| CC-8977 | 28.56 | 14.66 | 212.45 |
| STO-609 | 13.46 | 7.16 | 36.42 |

*Multi-drug-resistant patient.

**Supplementary Table S5. List of primers used for qRT-PCR gene expression analysis.**

| Gene | Forward [5'-3'] | Reverse [5'-3'] |
|---|---|---|
| *CaMKK2* | AGC TGA GGA CTT GAA GGA CCT GAT | AGG TTG TCT TCG CTG CCT TGC TT |
| *IL6* | AAC CTG AAC CTT CCA AAG ATG G | TCT GGC TTG TTC CTC ACT ACT |
| *CXCL10* | GTG GCA TTC AAG GAG TAC CTC | GCC TTC GAT TCT GGA TTC AGA CA |
| *APRIL* | CTC TGC TGA CCC AAC AAA CAG | TTT TCC GGG ATC TCT CCC CAT |
| *BAFF* | GGG AGC AGT CAC GCC TTA | CGT GGG AGG ATG GAA ACA CAC |
| *ACTNB* | CCT TGC ACA TGC CGG AG | GCA CAG AGC CTC GCC TT |

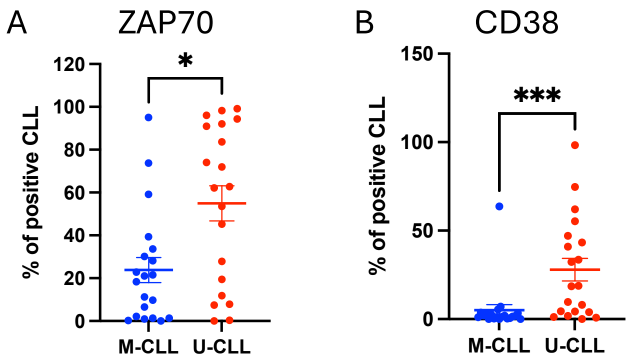


**Supplementary Figure S1. Canonical CLL prognostic markers CD38 and ZAP-70 are increased in IGHV-unmutated cases.** Flow cytometry was performed on peripheral blood $CD19^{+}$ CLL cells to quantify ZAP-70 (A) and CD38 (B) expression. Samples were stratified by IGHV mutation status (M-CLL vs U-CLL). Each symbol represents one patient; horizontal bars indicate mean ± SEM. Group differences were assessed using the Mann-Whitney test; significance is indicated by asterisks.

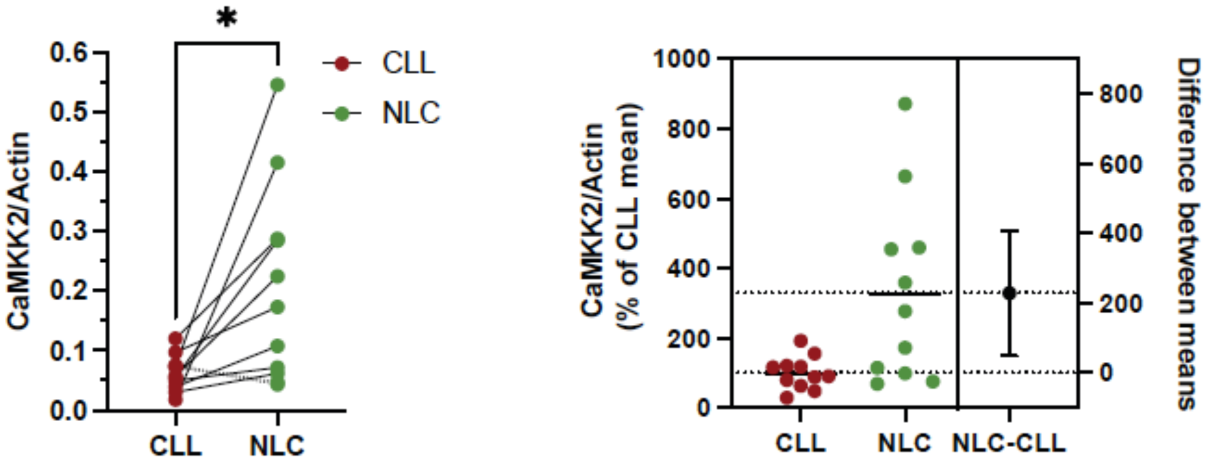


**Supplementary Figure S2.** ***CaMKK2*** **expression in CLL and corresponding autologous NLC.** Supplementary Figure S2. CaMKK2 expression in CLL cells and paired autologous NLC. PBMCs from CLL patients were cultured at high density for 14 days to generate a non-adherent $CD19^+$ CLL fraction and an adherent NLC-enriched fraction. Left: CaMKK2 mRNA expression (CaMKK2/ACTB) measured by qRT-PCR in purified $CD19^+$ CLL cells (CLL) and matched autologous adherent NLC; each line connects paired measurements from the same patient. Statistical significance was assessed using a two-tailed paired Wilcoxon signed-rank test; $p < 0.05$. Right: Estimation plot showing CaMKK2 expression in CLL and NLC, and the paired difference (NLC-CLL), expressed as a percentage of the mean CaMKK2/ACTB value in the CLL group; points represent individual patients and error bars [illegible]
